# Specific and versatile monoclonal antibodies for hantavirus research

**DOI:** 10.1101/2025.09.03.674135

**Authors:** Autumn LaPointe, Kimberly Martinez, Christina Shou, Inessa Manuelyan, Jason Botten, Alison M. Kell

## Abstract

Rodent-borne hantaviruses pose a continual public health threat to humans through zoonotic transmission, with case fatality rates of up to 50% in some cases. Human infections can lead to hemorrhagic fever with renal syndrome (HFRS) or hantavirus cardio-pulmonary syndrome (HCPS), depending on the viral species. Despite the morbidity and mortality associated with this family of viruses, no anti-viral therapeutics or vaccines are available to treat and prevent hantavirus disease. The relative shortage of commercially available reagents to study hantavirus infections *in vitro* and *in vivo* likely contributes to the challenges in developing viral countermeasures. This report describes the generation of a panel of mouse monoclonal antibodies that collectively recognize the four viral proteins of Seoul virus (*Orthohantavirus seoulense*), an Old World hantavirus with worldwide distribution and the causative agent of HFRS. We have validated the specificity and versatility of these antibodies against a subset of Old World and New World hantaviruses in assays relying on antigen recognition in denatured or native conformations. We present several antibodies that specifically recognize the Seoul virus nucleoprotein and polymerase protein in Western blotting and immunostaining assays. We also identified three novel antibodies directed against the glycoprotein complex that are capable of binding to the N-terminal glycoprotein of all hantaviruses tested. These antibodies are freely available to all hantavirus researchers to add to the small, but growing, collection of reliable and available reagents to be used to study hantavirus biology, identify novel antiviral compounds, and measure viral prevalence in the laboratory and the field.

**Importance:** Pathogenic hantaviruses cause severe hemorrhagic disease and pose a significant public health threat world-wide. Insufficient research into the biology of these viruses has slowed the development of effective direct-acting antivirals and vaccines. Here, we describe the generation and validation of novel, specific monoclonal antibodies for the detection of Seoul virus proteins *in vitro*. These reagents can be used to fill in critical gaps in knowledge regarding hantavirus entry, protein expression, and particle generation.

## Introduction

Hantaviruses (genus *Orthohantavirus*, order *Elliovirales*, class *Bunyaviricetes*) are zoonotic segmented, negative-sense RNA viruses that can cause severe hemorrhagic disease in humans. Old World hantaviruses (OW) are distributed throughout Southeast Asia, Russia, and Europe and cause a form of hantavirus disease termed Hemorrhagic Fever with Renal Syndrome (HFRS). New World hantaviruses (NW) are found in rodent reservoir populations in the Americas and are responsible for Hantavirus Cardio-Pulmonary Syndrome (HCPS). Infections with OW hantaviruses are more common with high morbidity but lower case-fatality rates (1-15%) compared to NW hantavirus infections (30-45%). Despite the public health concern, there are no FDA approved vaccines or targeted therapeutics for hantavirus infections.

The type species, *Orthohantavirus hantanense* (Hantaan virus, HTNV), was first isolated in 1978, and NW hantaviruses were discovered in 1993 (1-4). Yet much of the basic biology of this family of viruses remains underexplored. This can be explained, in part, due to a scarcity of tractable tools and models to study hantavirus infections *in vitro* and *in vivo*, especially compared to other segmented viruses, such as influenza. We report here the generation of monoclonal antibodies targeting the OW *Orthohantavirus seoulense* (Seoul virus, SEOV) nucleoprotein, glycoprotein complex, and polymerase and demonstrate their use for Western blotting and immunostaining assays. We tested these antibodies for specificity against SEOV and other NW and OW hantaviruses, finding cross-reactivity only in a subset of anti-glycoprotein antibodies. Several of the antibodies generated can be used to detect denatured proteins in cell lysates as well as native proteins in immunoassays. These tools will be made available for the research community through public repositories and have widespread applicability to test antiviral efficacy, study pathogenesis in culture models, and to explore virus-host interactions for this understudied virus family.

## Results

### Antibody development

Through Genscript, we purchased custom-generated hybridomas from mice immunized against the four SEOV proteins: nucleoprotein (N), N-terminal glycoprotein (G_n_), C-terminal glycoprotein (G_c_), and polymerase (L). Mice were immunized with linear peptides from Gn (WRKKANQESANQNSC), Gc (CGDPGDVMGPKDKPF), or L (LEKKVIPDHPSGKTC). For N, the entire recombinant protein was synthesized as the target antigen for immunization. From the immunization scheme, we received five hybridomas per protein target, selected for antigen reactivity by peptide ELISA. Supernatants from the hybridomas were purified on protein A/G columns, concentrated, and tested for reactivity and specificity by Western blot and immunostaining assays, as described below.

### Nucleoprotein

The hantavirus nucleoprotein coats the viral RNA, plays a critical role in viral transcription and translation, and is abundantly expressed in infected cells. We tested antibodies from each of the five hybridomas generated against the SEOV nucleoprotein for reactivity by Western blot (WB), immunofluorescence (IFA), and focus-forming unit assay (FFU). To test for specificity against SEOV N, we transfected HEK293T cells with plasmids expressing the N protein from the OW hantaviruses SEOV and HTNV, as well as the NW hantaviruses *Orthohantavirus sinnombreense* (Sin Nombre, SNV) and *Orthohantavirus andesense* (Andes virus, ANDV). Although expression for HTNV N was low, all ectopically-expressed proteins were detected in the lysates by their HA tag but only SEOV N was detected by the monoclonal antibodies (Fig 1a). Next, we tested the specificity of these antibodies in immunofluorescence assays (IFA).

**Figure 1.**
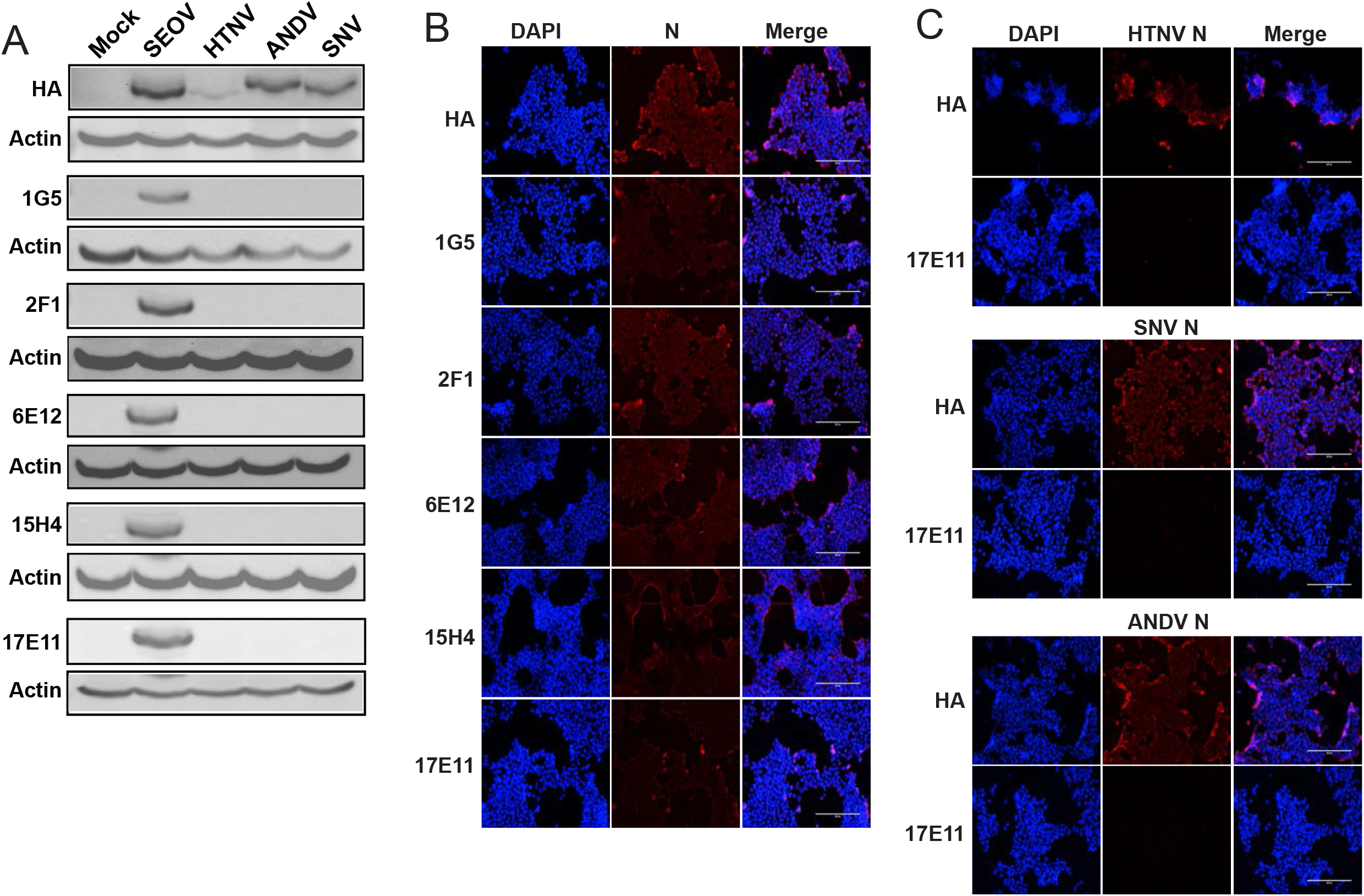
Antibodies specifically recognize SEOV N in transfected cells. HEK293T cells expressing HA-tagged N protein from SEOV, HTNV, SNV, and ANDV were interrogated by Western blotting and IFA. A) 30ug of cell lysates were loaded into 10% polyacrylamide gels in denaturing conditions and probed for N expression using anti-HA and newly generated anti-SEOV N antibodies. Actin served as a loading control. B) HEK293T cells transfected with SEOV N plasmid, probed using DAPI (nuclei, blue) and anti-HA or antibodies against SEOV N (red). C) HEK293T cells transfected with N plasmids from the indicated virus and probed with anti-HA antibodies or SEOV N antibody 17E11 (red) and DAPI (nuclei, blue). Cells visualized on the EVOS Cell Imaging System.

HEK293T cells were transfected with the N expression plasmids used above for SEOV, HTNV, SNV, and ANDV. Two days post-transfection, the cells were fixed, permeabilized, and probed with the monoclonal antibodies followed by an anti-mouse IgG fluorescent secondary. We observed that all transfected N proteins were detectable with the anti-HA primary antibody, but only SEOV N was detected with the anti-N monoclonal antibodies (Fig 1b, c). Fig 1c shows the absence of N staining for HTNV, SNV, and ANDV N using a representative anti-N, clone 17E11. Importantly, very low background staining was detected in mock transfected cells, supporting the utility of these antibodies for IFA (Supp Fig 1a).

We then asked whether these antibodies could detect N during infection. We infected Vero E6 cells with SEOV, HTNV, SNV, or ANDV for 12 days and performed WB on collected cell lysates (Fig 2a). Our standard hantavirus propagation protocols call for 12-day growth in Vero E6 cells as the peak of viral production. Therefore, this time point was chosen to be reasonably assured of abundant viral protein in cell lysates for all hantaviruses tested. Again, we see that the SEOV N antibodies are highly specific for SEOV N and can be used to assess SEOV infection status in susceptible cells. Similar results were observed when probing hantavirus-infected Vero E6 cells for IFA (Fig 2b). Again, we noted very low background staining for these antibodies on mock-infected cells (Supp Fig 1b). Finally, we tested the five anti-N antibodies for their efficacy in FFU assays. Hantaviruses are typically non-cytopathic in healthy cells and therefore do not form plaques. Instead, virus titrations can be executed by immunostaining. Vero E6 cells were infected with serial dilutions of SEOV for 1 hour, after which a methylcellulose overlay was added, and cells were incubated for 7 days post-infection. Cells stained with each of the anti-N antibodies showed well-defined foci of infection with very low background staining of uninfected monolayers (Fig 2c). These monoclonal antibodies, therefore, are specific for the SEOV nucleoprotein and are effective in denaturing WB conditions and IF assays.

**Figure 2.**
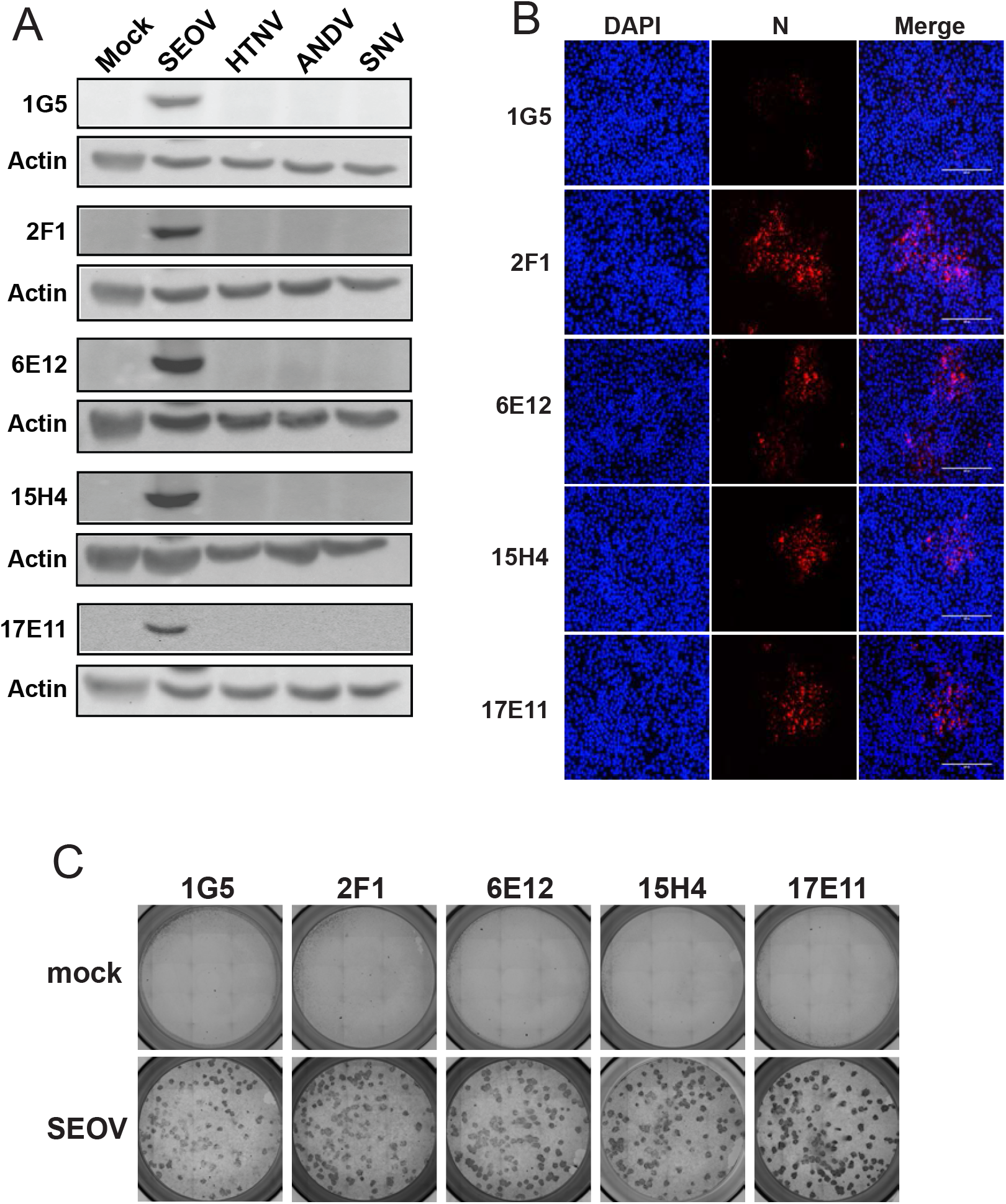
Antibodies specifically recognize SEOV N in hantavirus infected cells. Vero E6 cells were infected with SEOV, HTNV, ANDV or SNV and interrogated by Western blotting (A), IFA (B), and FFU assay (C). A) 30ug cell lysates were loaded into 10% polyacrylamide gels in denaturing conditions and probed for N expression using antibody clones against N. Actin serves as loading control. B) SEOV-infected Vero E6 cells probed with anti-N antibodies (red) and DAPI (nuclei, blue). Cells visualized on the EVOS Cell Imaging System. C) Focus forming unit assay using Vero E6 cells infected with SEOV and incubated with a 2% methylcellulose overlay for 7 days. Cells were probed with anti-N antibody clones, HRP-conjugated secondary antibody, and visualized for foci of stained cells, representing viral infectious units. Mock-infected cells represent non-specific background antibody interactions.

### Glycoproteins

The hantavirus M segment encodes a precursor glycoprotein that is co-translationally cleaved at a conserved WAASA site into two mature glycoproteins (G_N_ and G_C_) (5). These glycoproteins form trimeric complexes in the ER consisting of a G_N_ dimer and a G_C_ monomer (known as glycoprotein complex, GPC), and are trafficked to the Golgi (6-11). Co-expression of both proteins is necessary for trafficking from the ER to the Golgi, as individual expression results in retention of the proteins in the ER (12-17). We immunized mice with peptides from either G_N_ or G_C_, but found that none of the antibodies raised against G_C_-immunization were able to detect glycoproteins from any of the viruses tested either in transfected or virus-infected cells (data not shown). For these experiments, we transfected HEK293T cells with plasmids expressing the M segment sequence for both G_N_ and G_C_ (GPC) for SEOV, HTNV, and ANDV. These plasmids encode for the M segment coding region with an HA tag at the C-terminus. Thus, upon expression and cleavage of the polyprotein, only the 52-58kDa G_C_ protein will be detectable with an anti-HA antibody. SNV GPC was omitted because this plasmid failed to express detectable HA-tagged G_C_. Curiously, we found that several anti-G_N_ antibodies detected a protein migrating at the same size as our viral G_N_ (∼72-74kDa) in transfected HEK293T cells (Fig 3a). This cross-reactive protein was only detected in HEK293T and not in other commonly used cells for hantavirus research (A549, Vero E6, HUVEC-C, and primary rat lung microvascular endothelial cells) (Fig 3b). Surprisingly, we found that three of our tested anti-G_N_ antibodies (1B12, 11A5, and 12D10) detected the G_N_ from HTNV, SNV, and ANDV-infected Vero E6 cell lysates by WB (Fig 3c). Unfortunately, none were able to recognize native GPC by IFA (data not shown).

**Figure 3.**
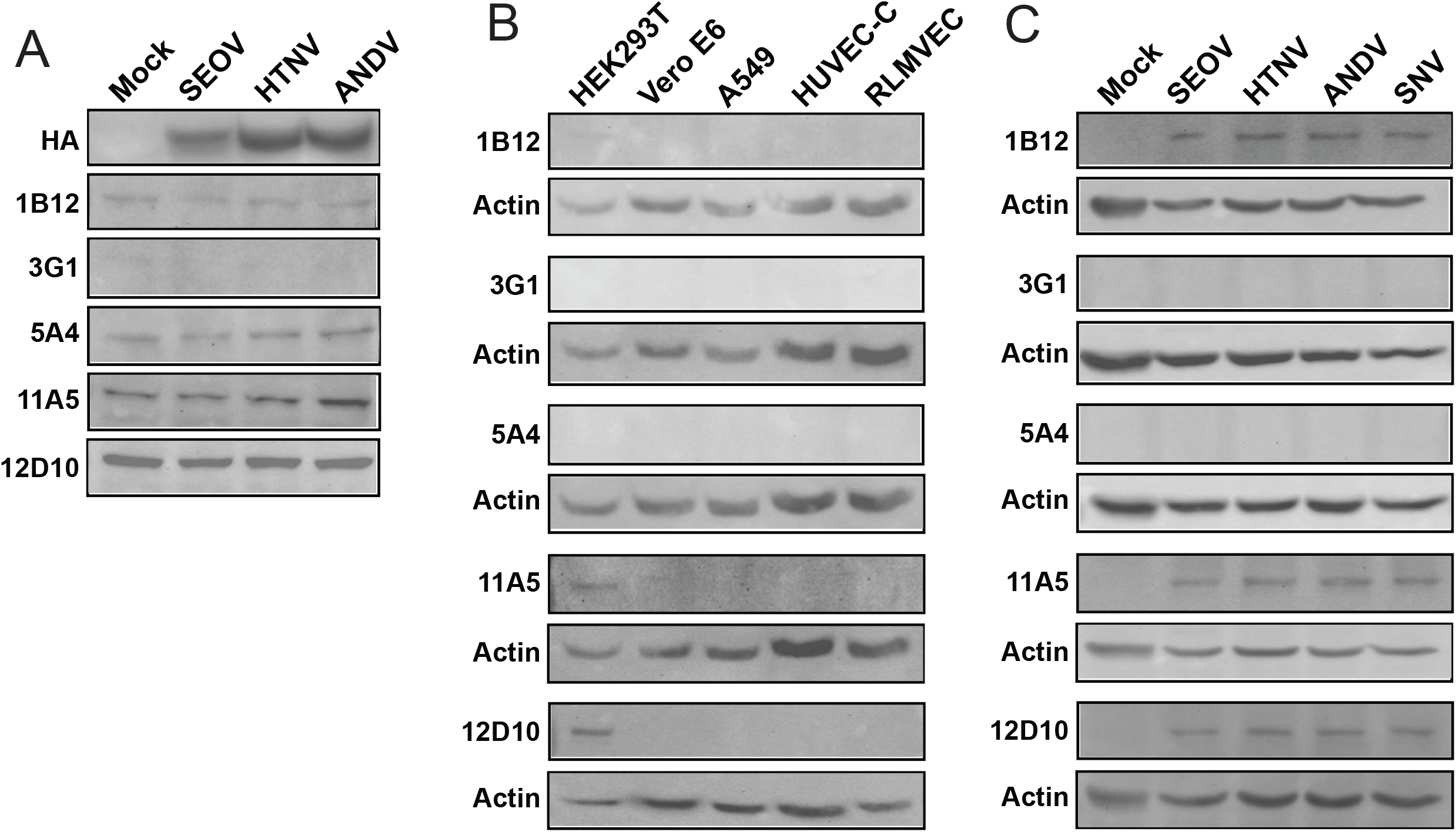
Several anti-G_N_ antibodies recognize G_N_ from OW and NW hantaviruses. A) HEK293T cells were transfected with plasmids expressing the indicated hantavirus GPCs from SEOV, HTNV, or ANDV. Each GPC encodes a C-terminal HA tag which, upon cleavage allows for the detection of expressed G_C_ only (52-58 kDa). Cell lysates were interrogated by Western blotting for recognition by anti-G_N_ antibodies or the anti-HA antibody. B) Un-transfected cell lysates from several cell types commonly used in hantavirus research were interrogated by Western blotting for cross-reactivity with anti-G_N_ antibodies. C) Vero E6 cells infected with SEOV, HTNV, ANDV, SNV, or mock infected were analyzed by Western blot for detection of G_N_ by anti-G_N_ monoclonal antibodies. Actin served as a loading control.

### Polymerase

The RNA-dependent RNA polymerase protein of hantaviruses (L) is expressed at low, almost undetectable levels in cultured cells due to its presumed toxicity (18, 19). Even overexpression of plasmid derived L protein has proven difficult without mutations that inhibit its transcriptional activity (18). We generated HA-tagged plasmids for ectopic expression of SEOV L and SNV L in HEK293T cells to test the anti-L antibodies in WB and IF assays (Fig 4a, b). Two of the four antibodies tested were able to detect SEOV L by both assays using transfected HEK293T cells, but none were cross-reactive against SNV L. We also fund that clone 13G8 was able to detect SEOV L from infected Vero E6 cell lysates, but not from cells infected with HTNV, SNV, or ANDV (Fig 4c). Despite considerable effort, we were unable to detect L expression in SEOV-infected cells by IFA (data not shown).

**Figure 4.**
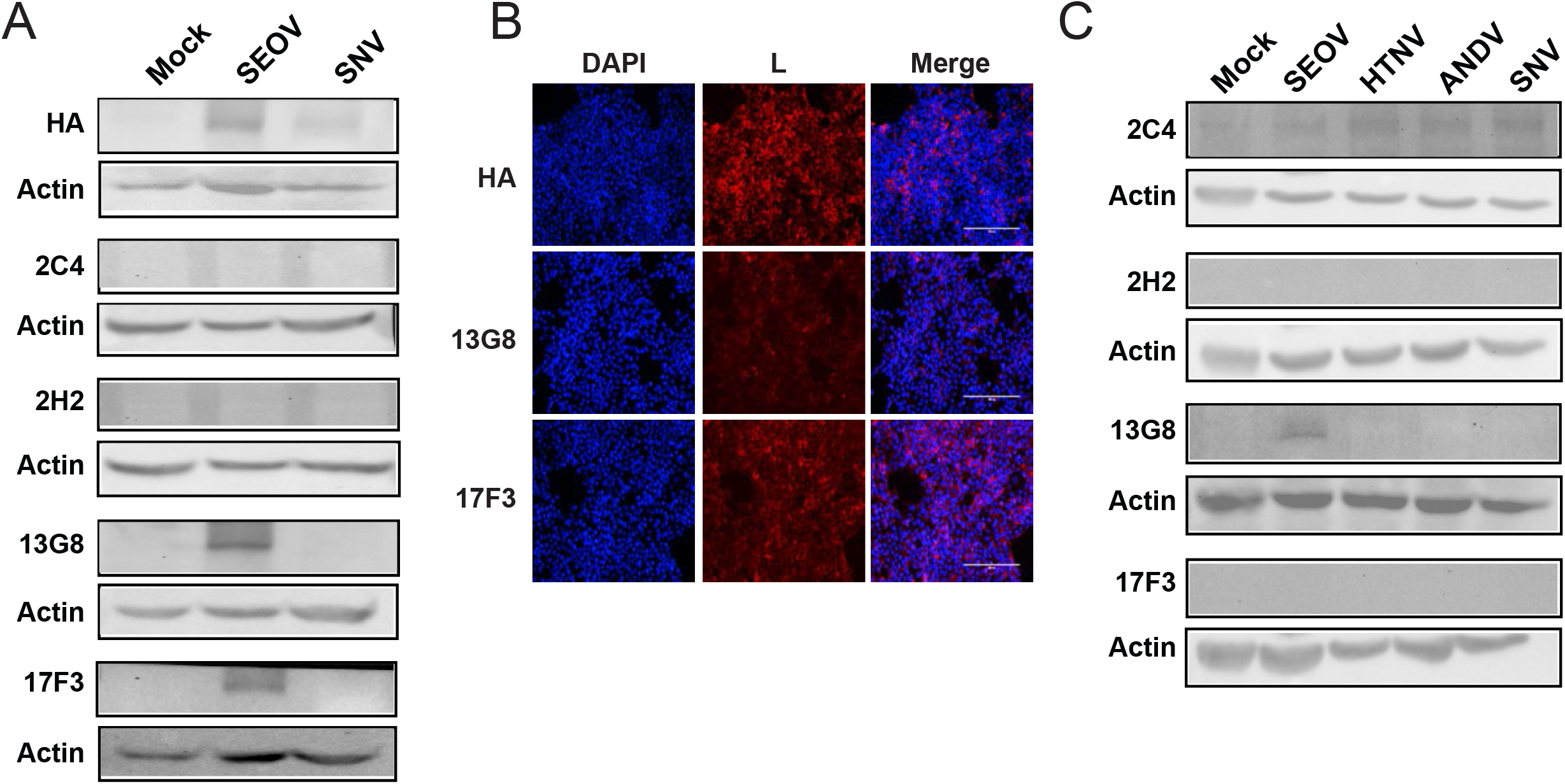
Monoclonal antibodies specifically recognize SEOV L protein. A) HEK293T cells transfected with plasmids expressing HA-tagged SEOV L or SNV L were analyzed by Western blotting using anti-L monoclonal antibodies. B) SEOV L plasmid transfected HEK293T cells were probed with anti-HA antibody or clones 13G8 or 17F3 (red) and DAPI (nuclei, blue). Cells visualized on the EVOS Cell Imaging System. C) Vero E6 cells infected with SEOV, HTNV, ANDV, SNV, or mock-infected were analyzed by Western blot for detection by anti-L monoclonal antibodies. Actin served as a loading control.

## Conclusions

We have characterized novel monoclonal antibodies for their specificity and versatility for detection of hantavirus proteins in exogenous and endogenously expressed conditions. These tools for SEOV detection and quantification should replace less specific and less reliable commercial sources for hantavirus antibodies. We are actively working to make these reagents available to the hantavirus research community though trusted biorepositories, such as BEI Resources.

## Methods

### Cell culture

Vero E6 cells (ATCC, CRL-1586), A549 (ATCC, CCL-185), and HEK293T (ATCC, CRL-3216) cells were cultured in Dulbecco’s modified Eagle’s medium (DMEM) supplemented with 10% heat-inactivated FBS, 1% pen/strep, 1% nonessential amino acids, 2.5% HEPES. Primary rat microvascular endothelial cells (RLMVEC, VEC Technologies) were cultured in MCDB-131 base medium (Corning) supplemented with EGMTM-2 Endothelial SingleQuots Bullet Kit (Lonza, CC-4176) and 10% heat-inactivated FBS. Human umbilical endothelial cells (HUVEC-C; ATCC, CRL-1730) were cultured in Lonza EGM-Plus (Lonza, CC-4542) supplemented with bullet kit (Lonza, CC-5036) and 10% heat-inactivated FBS. All cells were cultured in tissue culture-treated plastics and plate for endothelial cells were coated with rat-tail collagen (VWR, 47747-218).

### Viruses and *in vitro* infections

Seoul virus (strain SR11), Hantaan virus (strain 76-118) Sin Nombre virus (strain NMR11), and Andes virus (CHI7913) were propagated on Vero E6 cells (ATCC, CRL-1586) for 12 days, with a maximum of 3 passages. Infectious virus was isolated by harvesting supernatant and centrifuging at 1000 rpm for 10 minutes to remove cellular debris. For virus infections, cells were seeded in cell culture vessels 18-24 hours prior to infection at a target density of 70%. Virus stock was diluted to the target MOI using serum-free Dulbecco’s modified Eagle’s medium (DMEM) supplemented with 1x pen/strep, 1% nonessential amino acids, and 2.5% HEPES. Virus was allowed to adsorb for 1 hour at 37ºC. Cells were washed twice with sterile PBS solution (FisherScientific, SH30264FS), and appropriate culture medium was added for the duration of the experiment.

### Antibody preparation

Hybridomas were cultured in Dulbecco’s modified Eagle’s medium (DMEM) supplemented with 10% heat-inactivated FBS, 1% pen/strep, 1% nonessential amino acids, 1% L-Glutamine, 2.5% HEPES, and 1x hybridoma fusion and cloning supplement (HFCS; MilliporeSigma 11363735001) for the first passage. Hybridomas were then maintained in in Dulbecco’s modified Eagle’s medium (DMEM) supplemented with 10% heat-inactivated FBS, 1% pen/strep, 1% nonessential amino acids, 1% L-Glutamine, and 2.5% HEPES. Supernatants were collected every 2-3 days when cells were at 90% confluency. They were then centrifuged at 1,000xg for 10min to remove cell debris and stored at 4C until purified. To purify antibody, a 2mL Pierce protein A/G agarose column (Thermo Scientific, 20422, 29922) was prepared according to manufacturer instructions. Supernatants from the same hybridoma culture were pooled together and 1/3 volume of Pierce protein A/G IgG binding buffer (Thermo Scientific, 54200) was added to the supernatants. Supernatant and binding buffer were run through the column according to manufacturer instructions. Antibody was eluted from the column using 8mL of Pierce IgG elution buffer (Thermo Scientific, 21004) and neutralized with 400uL 1M Tris pH 7.5. Amicon Ultra-15 centrifugal filters, 30kDa MW (MilliporeSigma, UFC903024), were used to concentrate the purified antibody down to 200uL. Antibody concentration was determined via Pierce BCA Protein Assay Kit (ThermoFisher Scientific, 23225) and stocks stored at 1ug/mL.

### Plasmids

To validate the detection of hantavirus proteins by each monoclonal antibody, we subcloned each of the described proteins from SEOV, HTNV, SNV, and ANDV into our previously described pCAGGS expression vector (20, 21). This vector contains the coding sequence of each viral protein followed by a C-terminal hemagglutinin (HA) epitope tag (YPYDVPDYA). Viral protein coding sequences were PCR amplified from viral cDNA and subcloned into the vector using Gateway Technology (Invitrogen) following manufacturer’s instructions. The nucleotide sequence of each plasmid was verified by DNA sequencing (Plasmidsaurus). Viral protein sequences were subcloned from the following virus strains: SEOV strain SR-11, HTNV strain 76-118, SNV strain NMR11, and ANDV strain CHI-7913. For the SEOV L protein plasmid, the previously reported K44A mutation was made to increase expression of L (18).

### Plasmid transfections

HEK 293T cells were seeded in 10cm dishes at 2×10^5 cells/mL. Transfections were accomplished 24hrs later using jetPRIME (VWR, 89129-924) according to manufacturer’s specifications. Briefly, 5ug (GPC and L) or 10ug (N) of the indicated expression plasmid were diluted in 500uL jetPRIME buffer and 10uL of jetPRIME reagent was added. The mixture was incubated at room temperature for 10min and was then added dropwise to the 10cm dishes of cells. Lysates were harvested 48hrs post-transfection in RIPA buffer (50mM Tris HCl pH 7.4, 150mM NaCl, 1% (v/v) NP-40, 0.5% (w/v) Na-deoxycholate) and clarified through 25,000rcf centrifugation for 15 minutes at 4ºC. Protein was quantified via Pierce BCA Protein Assay Kit (ThermoFisher Scientific, 23225). 30ug of protein was loaded per sample into an 8% (L) or 10% (N and GPC) polyacrylamide gel and, after denaturing electrophoresis, transferred to a 0.45µm nitrocellulose membrane (VWR, 10120-006). Membranes were blocked at room temperature in 10% FCS in PBS-T. Primary antibodies (1ug/mL) were incubated at 1:500 at 4º overnight. HRP-conjugated secondary antibodies against primary antibody species were incubated 1:10,000 for 1 hour at room temperature. Blots were imaged on BioRad Chemidoc MP Imaging System using chemiluminescence Pierce Substrate for Western Blotting (VWR, PI80196).

### Generation of infected-cell lysates

Cell lysates to be interrogated for protein expression by immunoblot analysis were harvested in protein lysis buffer 12 days post-infection and prepared as previously described (22). Briefly, protein was harvested in RIPA buffer (50mM Tris HCl pH 7.4, 150mM NaCl, 1% (v/v) NP-40, 0.5% (w/v) Na-deoxycholate) and clarified through 25,000rcf centrifugation for 15 minutes at 4ºC. Protein was quantified via Pierce BCA Protein Assay Kit (ThermoFisher Scientific, 23225). 30ug of protein was loaded per sample into an 8% (L) or 10% (N and GPC) polyacrylamide gel and, after denaturing electrophoresis, transferred to a 0.45µm nitrocellulose membrane (VWR, 10120-006). Membranes were blocked at room temperature in 10% FCS in PBS-T. Primary antibodies (1ug/mL) were incubated at 1:500 at 4º overnight. HRP-conjugated secondary antibodies against primary antibody species were incubated 1:10,000 for 1 hour at room temperature. Blots were imaged on BioRad Chemidoc MP Imaging System using chemiluminescence Pierce Substrate for Western Blotting (VWR, PI80196).

### Immunofluorescence assay

Vero E6 cells were seeded in 48 well plates at 2.5×10^3 cells/well. When cells had reached 70-80% confluency, they were infected with either SEOV, HTNV, ANDV, or SNV at an MOI of 0.1. Virus was allowed to adsorb for 1hr at 37^°^ C, then cells were washed 2x with 1x PBS and complete DMEM was added. At 4 days post-infection, supernatants were removed and cells were washed 1x with 1x PBS. Cells were fixed with 95% ethanol:5% acetic acid for 10min at - 20^°^ C. Cells were permeabilized with 0.01% Triton X-100 in PBS for 15min at room temperature and then blocked with 3% FBS in PBS for at least 30min. Primary antibodies (1ug/mL) were incubated at 1:500 at 4^°^C overnight. Goat α mouse 647 fluorescent secondary antibodies (Invitrogen A21235) were incubated at 1:400 for at least 2hrs at room temperature. Nuclei were stained with DAPI at 1:1000 for 5min at room temperature. Stained cells were visualized using the 20x objective on the EVOS Cell Imaging System (ThermoScientific). Microscope settings for fluorescence intensity were set on mock-transfected/infected cells to remove background and remained constant for all imaging of transfected/infected cells.

### Focus Forming Unit Assay

Infectious virus was quantified using immunostaining as previously described (23). Vero E6 cells were infected with 100μL of culture supernatants and incubated for 2 hours at 37ºC. After incubation, a 2% carboxymethylcellulose overlay containing supplemented DMEM (2% pen/strep, 2% nonessential amino acids, 2% HEPES, 4% heat-inactivated FBS). Cells were incubated for 7 days at 37ºC. After incubation, cells were fixed with 95% EtOH:5% Acetic Acid for 10 minutes at -20ºC, blocked with 3% FBS in PBS, and probed for SEOV N (α SEOV nucleocapsid, 17E11 or others presented here) or HTNV N (anti-HTNV nucleocapsid 76–118, BEI resources NR-12152, cross-reacts with HTNV, SNV, and ANDV) at 4º for 24 hours. HRP conjugated secondary (donkey α rabbit for α HTNV, Jackson Immunoresearch 711-035-152; or goat α mouse for α SEOV, Jackson Immunoresearch 115-035-003) for 2 hours at room temperature. Foci were then stained using Vector VIP Substrate Kit (Vector Laboratories, SK-4600) and counted under a light microscope to calculate titer.

## Acknowledgments

Work performed by KM and CS was supported by the National Institutes of Health NIAID grant 1R21AI175721-01A1 granted to AMK and JB. ATL was supported by a post-doctoral fellowship by the National Institutes of Health NIGMS T32AI007538. The funders had no role in study design, data collection and analysis, decision to publish, or preparation of the manuscript. The authors also thank Dr. Rebekah Honce for technical support in plasmid preparation and expression.

---

**Supplementary Figure 1.**
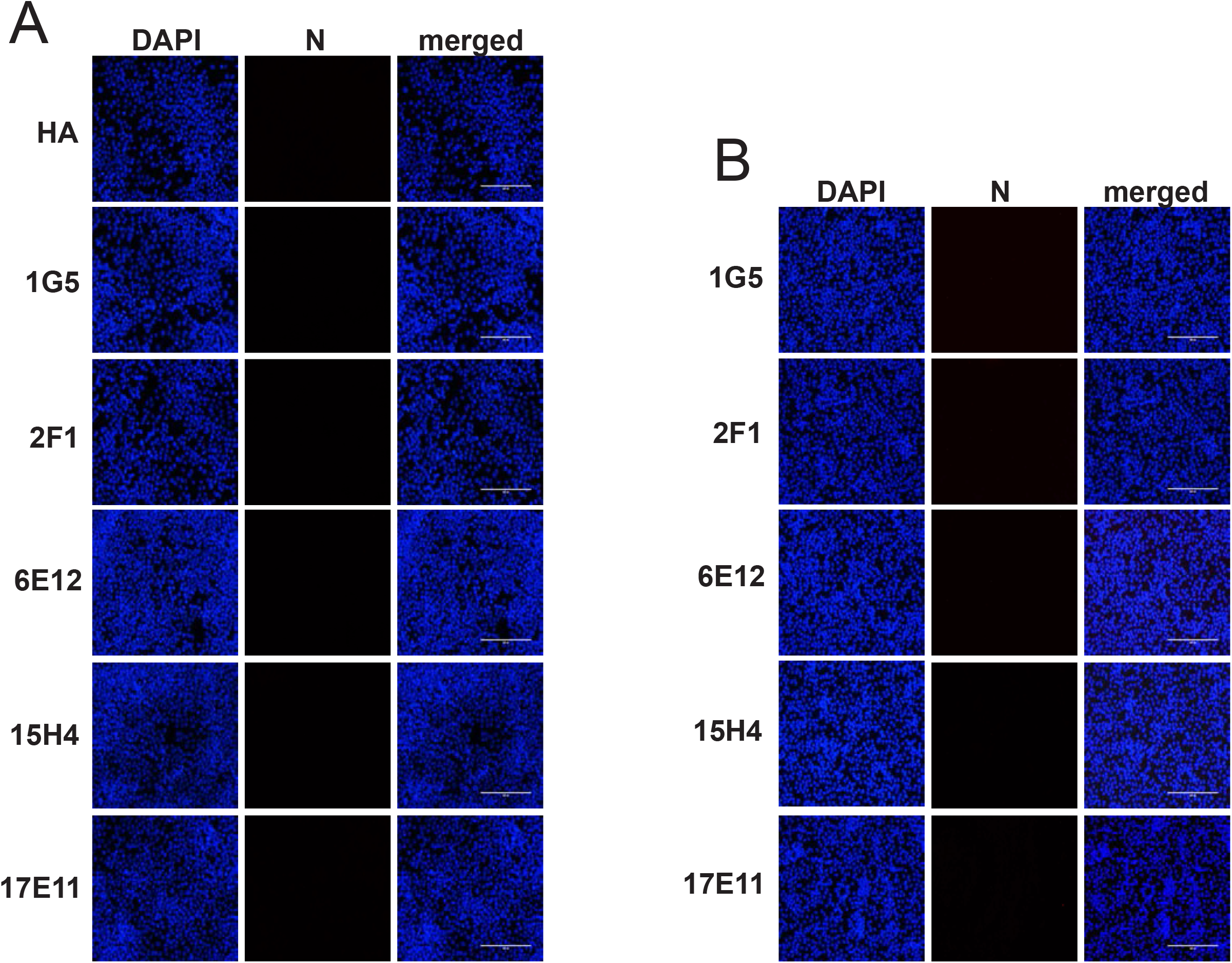
Anti-N antibodies exhibit low background staining in mock transfected or mock infected cells. Mock-transfected HEK293T cells (A) and mock-infected Vero E6 cells (B) probed with anti-HA and anti-N monoclonal antibodies (red) and DAPI (nuclei, blue). Cells visualized on the EVOS Cell Imaging System.

**Supplementary Figure 2.**
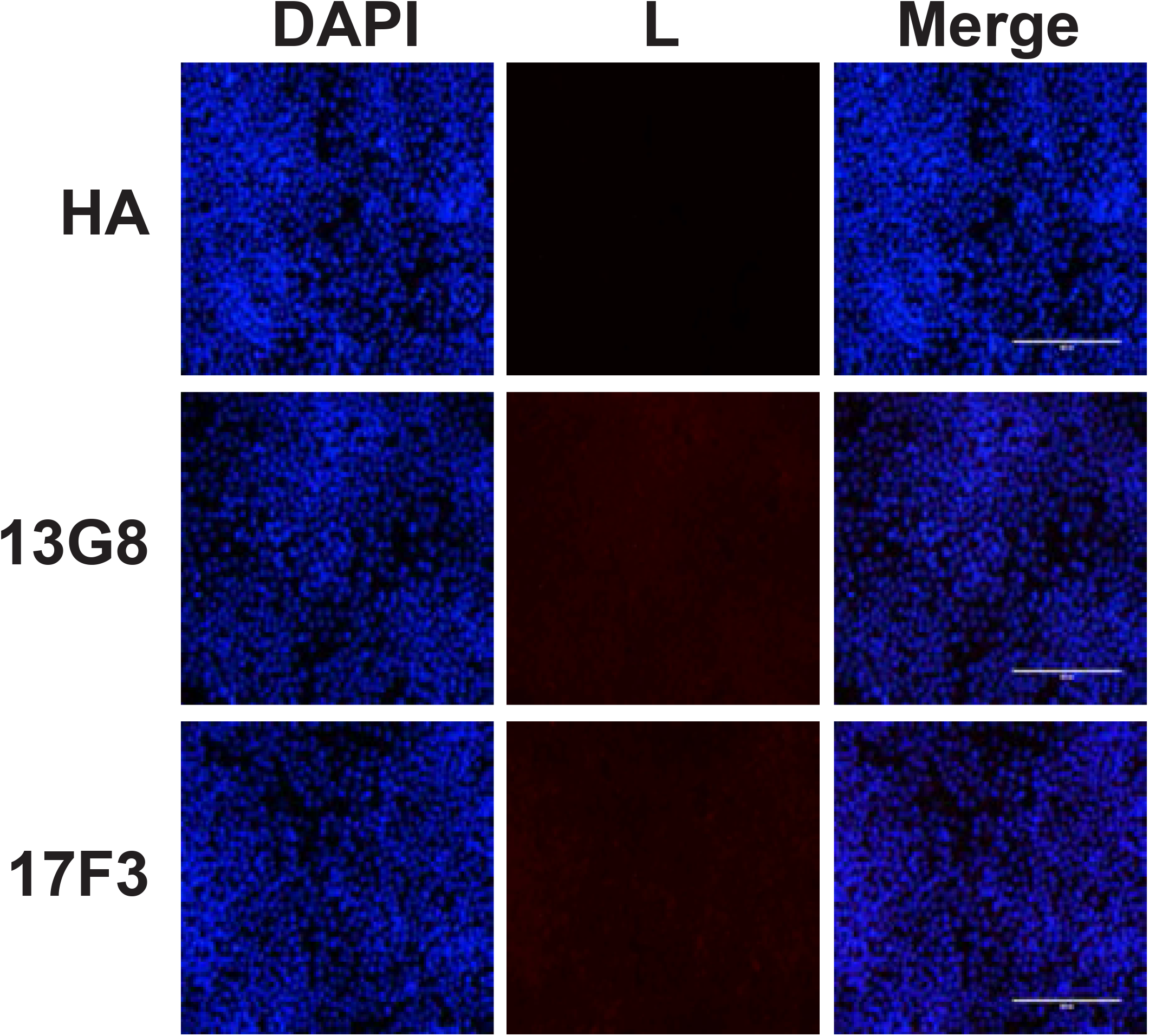
Anti-L antibodies exhibit low background staining in mock transfected cells. Mock-transfected HEK293T cells probed with anti-HA and anti-L monoclonal antibodies (red) and DAPI (nuclei, blue). Cells visualized on the EVOS Cell Imaging System.

